# G-quadruplex secondary structure from circular dichroism spectroscopy

**DOI:** 10.1101/184390

**Authors:** Rafael del Villar-Guerra, John O. Trent, Jonathan B. Chaires

**Affiliations:** James Graham Brown Cancer Center, University of Louisville, 505 S. Hancock St., Louisville, KY 40202 USA

## Abstract

A curated library of circular dichroism spectra of 23 G-quadruplexes of known structure was built and analyzed. The goal of this study was to use this reference library to develop an algorithm to derive quantitative estimates of the secondary structure content of quadruplexes from their experimental CD spectra. Principle component analysis and singular value decomposition were used to characterize the reference spectral library. CD spectra were successfully fit to obtain estimates of the amounts of base steps in *anti-anti, syn-anti* or *anti-syn* conformations, in diagonal or lateral loops or in other conformations. The results show that CD spectra of nucleic acids can be analyzed to obtain quantitative structural information about secondary structure content in an analogous way to methods used to analyze protein CD spectra.

---

Circular dichroism (CD) spectroscopy is a primary tool for the characterization of G-quadruplex (G4) structures. G-quadruplexes are functionally important genomic elements that form at specific locations in an orchestrated manner throughout the cell cycle.^[1]^ Different G4 structures, arising from differences in G-quartet stacking, strand segment orientation and loop arrangements, display unique CD spectral signatures.^[2]^ Qualitative rules-of-thumb have evolved that associate CD spectral features with particular G4 topologies, namely parallel (≈264 nm max, ≈245 nm min), antiparallel (≈295 max, ≈260 min) or “hybrid” (or 3+1) (≈295 max, ≈260 max, ≈245 min). ^[3] [4] [5] [6] [7]^ While some exceptions to these rules have been noted, they are generally accepted for the characterization and validation of quadruplex formation in potential quadruplex forming sequences.

CD spectroscopy is widely used for the quantitative determination of the secondary structural content of proteins. Over several decades, reference libraries assembled for this purpose have grown in content and a number of analytical algorithms have evolved that now make the quantitative analysis of protein spectra by CD fairly routine. Such is not the case for nucleic acids. For duplex DNA, CD is used primarily in a qualitative way to distinguish common secondary structures *(e.g.,* B-, A- and Z-forms) and is particularly valuable for monitoring changes in secondary structure in titration, binding or thermal denaturation experiments.^[2]^ A recent chemometric analysis of nucleic acid CD spectra was used to classify nucleic acid structures using a library of sequences and structures that expanded the range of topologies to include multistranded triplex and quadruplex forms.^[8]^ As of yet no quantitative analysis has been attempted for nucleic acids that is analogous to the approach used to quantify CD spectra of proteins to obtain more detailed structural information. The goal of this study is to develop such an analytical approach to provide quantitative secondary structural information about quadruplexes from their measured CD spectra.

A curated library of 23 CD spectra was built using sequences for which high-resolution structures were reported and deposited in accessible databases. The library is shown in Table S1 (supporting information). CD spectra were measured for each sequence, after appropriate annealing and sample preparation, under solution conditions identical to those used in the original structural determination using published protocols developed in our laboratory.^[9]^ Sample homogeneity was confirmed by sedimentation velocity experiments as previously described. ^[10] [11]^ In only one case (sample 186D) was significant heterogeneity observed, requiring additional purification by HPLC methods developed and reported by our laboratory. ^[12]^

Figure 1A shows the CD spectra obtained for the quadruplex library (these spectra are provided in digital form as an Excel file in supporting information). Spectra are normalized to molar circular dichroism (Δε) using molar strand concentration as a reference. The color coding in Figure 1A emphasizes that spectra fall into three groups as is evident by inspection, but also as determined by an unbiased principle component analysis (PCA) as will be described.

**Figure 1.**
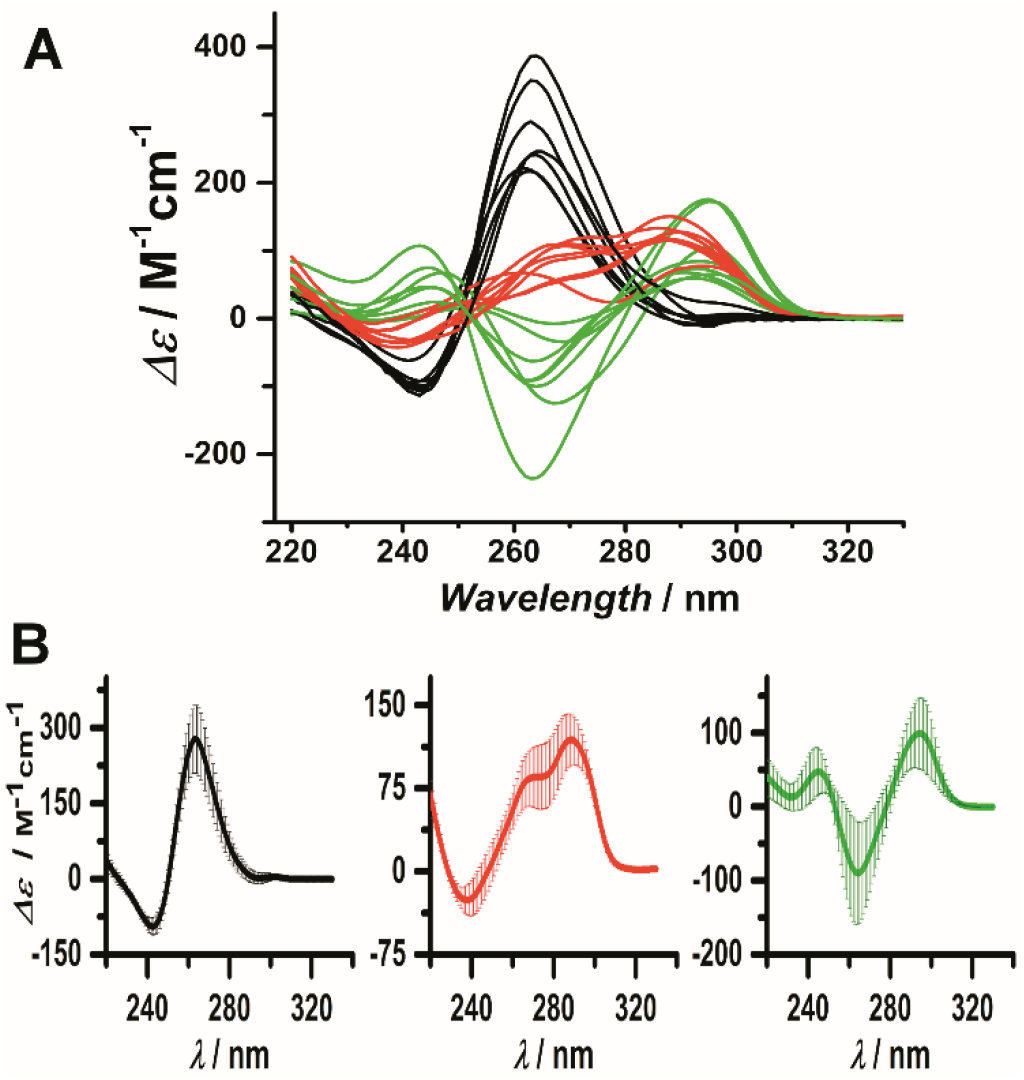
A) CD spectra of the reference library of 23 G-quadruplex. B) Average CD spectra obtained for cluster 1(left, black), cluster 2 (center, red) and cluster 3 (right, green). The associated standard deviations are represented by the shaded areas.

We used PCA and cluster analysis as an unbiased quantitative classification method to reduce any ambiguity in the assignment of a CD spectrum to a G-quadruplex conformational group to improve upon previously used semi-empirical and visual approaches. The results of PCA are shown in Figure 2A as a score plot in which the first two principle components are plotted against one another. Three clusters arise from the spectral data, as indicated by the shaded ellipses in Figure 2A. A dendrogram obtained by hierarchical clustering is shown in Figure 2B. Correlation of these clusters with the known topologies of the quadruplex structures within them reveals the separation of spectra into parallel (black), antiparallel (green) and “hybrid” or 3+1 (red) classes. The spectra in panel 1A are colored according to these classes. The average spectra for each class is shown as Figure 1B. Loading vectors are shown as blue lines in the score plot in Figure 2A. For the spectral data, these vectors show the most important and distinctive wavelengths that drive the PCA clustering: 258 nm for the parallel form, 234 nm for the antiparallel form and 284 nm for the “hybrid” form (Figure S2). These wavelengths supplement and extend the rules-of-thumb described above for the qualitative classification of quadruplex structures.

**Figure 2.**
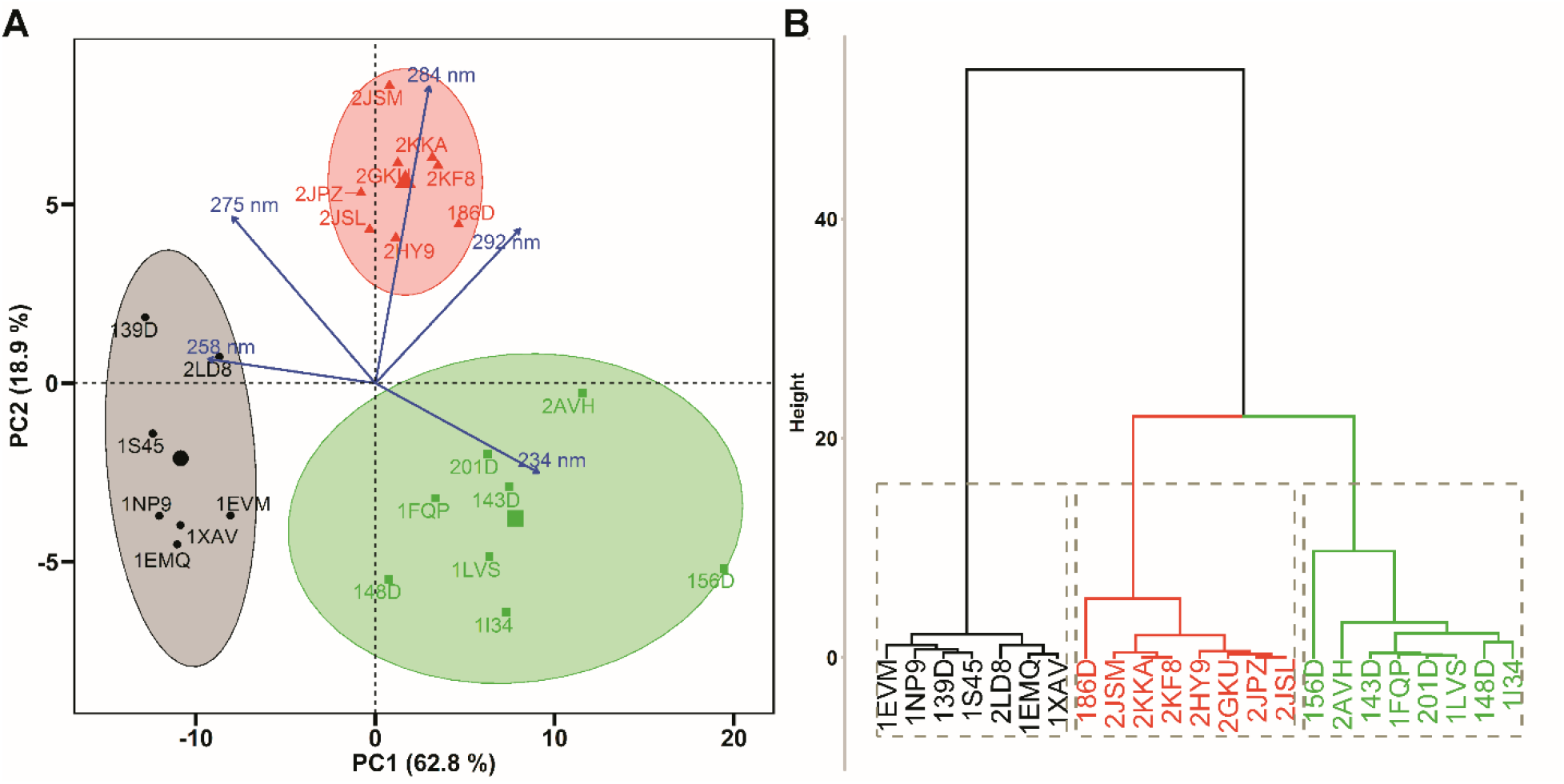
(A) Principal component analysis (PCA) and (B) Hierarchical Clustering on Principal Components (HCPC) of the reference CD spectra library of G-quadruplexes. In panel (A), the score plot is shown in which the first and second principal components are plotted against one another. The blue arrows indicate the loading vectors that drive the clustering.

This PCA and cluster analysis might be used in a number of ways. For analysis of a CD spectrum obtained for a new potential quadruplex forming sequence (properly normalized to Δε), an unbiased quantitative classification with respect to the reference spectra clusters would allow its most probable topology to be inferred. This could be easily done by adding the unknown spectrum to the reference library spectra and running the PCA to determine into which cluster the new spectrum falls. Alternatively, if the position of the new spectrum falls outside the clusters, this would provide unambiguous evidence that the spectrum arises from either a mixture of known quadruplex structures or from a structure not contained in our reference library. Finally, if the new spectrum is thought to be a mixture, the average spectra for the topological classes shown in Figure 1B could be used along with nonlinear least squares fitting to quantitatively estimate the composition of the mixture in terms of the reference quadruplex structures.

A quantitative secondary structural analysis of quadruplexes from their CD spectra is a more ambitious and complicated task. While it is well established that the percentage of secondary structural elements of proteins (e.g. α-helix, antiparallel β-sheet, parallel β-sheet, β- turn, polyproline II) can be obtained from their CD spectra,^[13]^ no analogous structural elements have yet been defined for quadruplexes. In order to develop such an analytical approach, it is necessary to define the secondary structural elements that make a significant contribution to the CD signal of a G-quadruplex, then to determine their underlying basis spectra by some analytical approach. For this purpose, we used singular value decomposition (SVD) and least square fitting to obtain structural information. This approach is analogous to that used for the structural determination of proteins by CD spectroscopy.^[14]^ The method assumes that the CD spectrum of a G-quadruplex can be represented as a linear combination of structural basis spectra

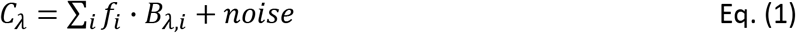

where *C_λ_* is the quadruplex CD spectra, f_*i*_ is the fraction of the *i* th structural element, and B_*λ,i*_ is the basis spectrum corresponding to the *i* th structural parameter. These fractions and basis spectra correspond to structural motifs that contribute to the G4 CD spectrum, and the first task is to determine what these are.

The number of structural elements that can be used is limited by the information content in the family of reference spectra.^[15] [14b, c]^ The results of SVD analysis of the reference spectra library are shown in supporting information (Figures S3-7 and Tables S2-3). We found that only five basis spectra were necessary to reconstruct the original CD spectra of our reference library within experimental error. Therefore, only five structural elements at most can be determined with confidence from our reference library.

Although any number of structural elements *(e.g.* loop types, glycosidic bond angles, quartet stacking geometry, strand orientations, etc.) might be used to describe and classify DNA quadruplexes topologies ^[6, 16]^, some of these may not make significant contributions to the CD signal. Since the CD of quadruplexes arises primarily from the stacking arrangements of guanine base steps within the G-quartet stacks, we have chosen the following approach to define five structural elements for each quadruplex. (Figure 3).

**Figure 3.**
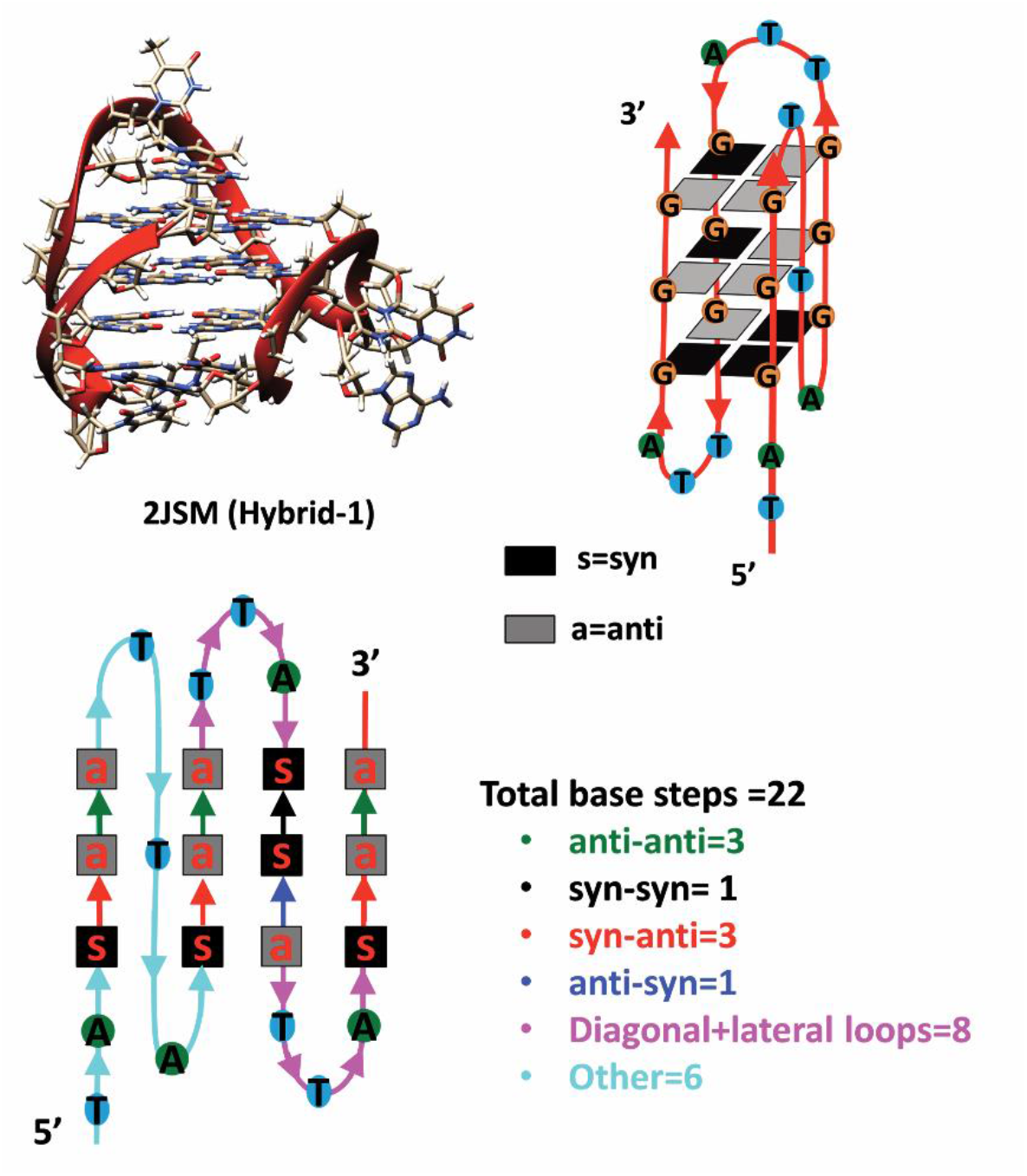
Schematic illustration of the definition of the G-quadruplex secondary structure elements used in this study. The example shown is for the human telomere hybrid 1 structure with the PDB identifier 2JSM.

Taking the total number of base steps in the strand of a particular structure as a reference, we define the fraction of each structural element as the number of base steps found in that element relative to the total strand length. We first count the guanine-guanine base steps in each of the glycosidic bond conformations *anti-anti, syn-anti* and *anti-syn* (see Figure 3). The progression from the 5’ to 3’ end of the first run of the G-quartets aligned according to the frame of reference were used to define the polarity of the G-G stacking base steps.^[16b]^ (Note that *syn-syn* guanine stacking is not counted because such steps rarely occur in the structures in our reference library or generally in any G4 structures.) Next, the fraction of base steps in either diagonal or lateral loops are counted. Finally, all remaining base steps are defined as “other”, and might include those in chain-reversal loops or terminal nucleotides that may or may not be stacked upon end quartets. Five structural elements are thus defined for each quadruplex, the fractions of which must sum to 1.0. The tabulated secondary structural element fractions for all members of our reference library are shown in Table S4 (supporting information).

The basis spectra of each chosen structural element were calculated by SVD analysis of the reference set and the matrix of fractions of structural elements, as described in more detail in the supporting information and shown in Figure S8. The shapes and signs of these basis spectra for dinucleotide guanine steps are generally consistent with the results of a simplified exciton coupling approach and more refined quantum mechanical calculations used to predict the CD spectra expected for different dinucleotide stacking orientations. ^[3, 6, 17]^ The basis spectrum obtained for lateral and diagonal loops are highly reminiscent of CD spectra obtained for single-stranded DNA di- and trinucleotide sequences. ^[18]^ The remaining basis spectrum for “other” has low amplitude and is nearly featureless.

These basis spectra can now be used to estimate the fraction of each structural element in a quadruplex of unknown structure by fitting its measured CD spectrum to eq 1. Figure 4 illustrates the procedure. Full details of the fitting procedure are described in supporting information.

**Figure 4.**
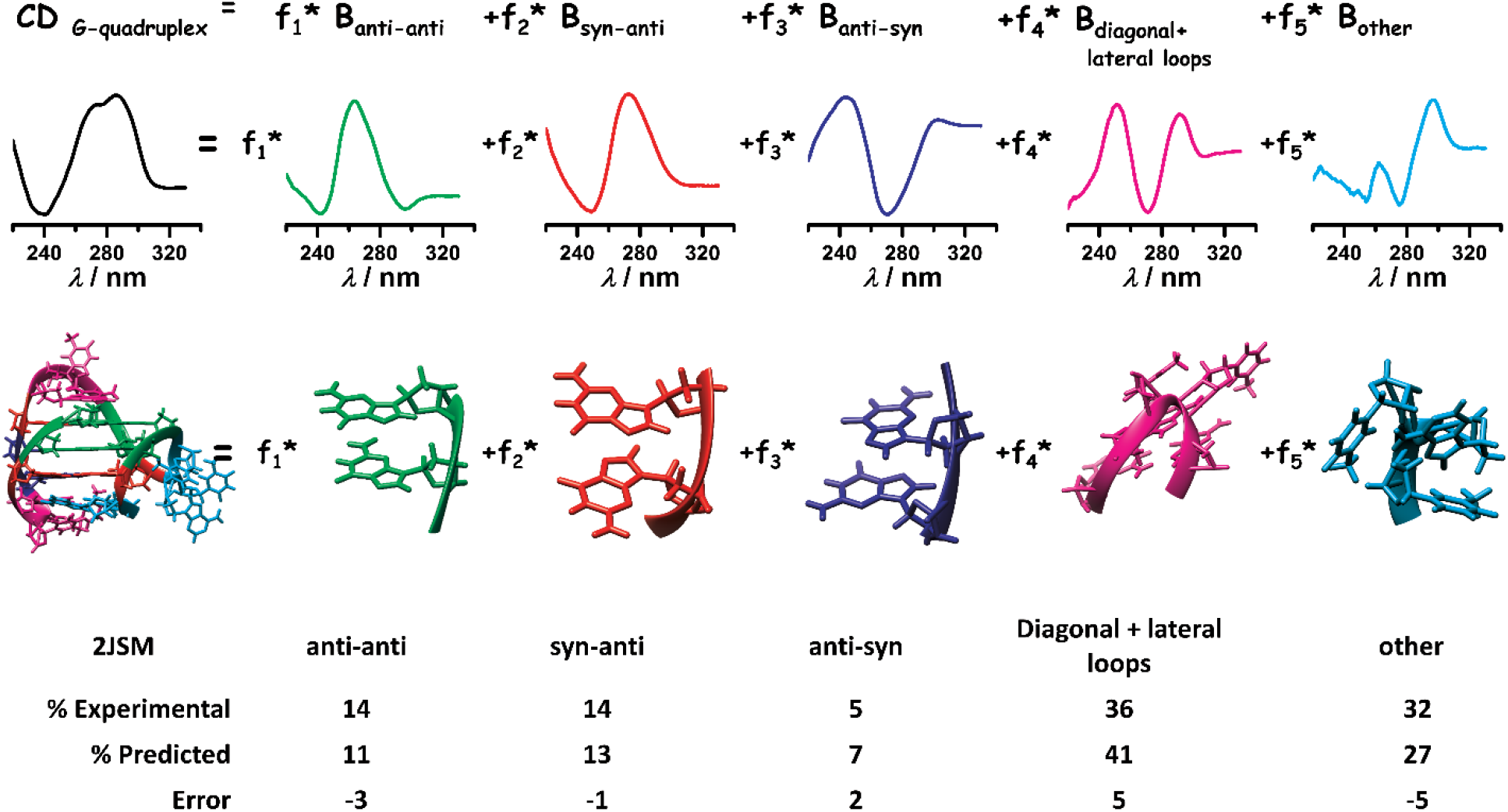
Schematic illustration of the method used to calculate the fractions of different secondary structural elements by constrained least-square fitting of a test CD spectrum to the five secondary basis spectra.

Figure 5 shows selected examples of fits of experimental CD spectra to obtain estimates of the fraction of *anti-anti, syn-anti, anti-syn* dinucleotide steps, lateral and diagonal loops, and other residual structures. Fits to all members of our reference library are shown in supporting information (Figure S9). The fits are remarkably good (small values of NRMSDspectral, Figure S10), and yield, for the first time, quantitative estimates of the fraction of structural elements in each quadruplex structure.

**Figure 5.**
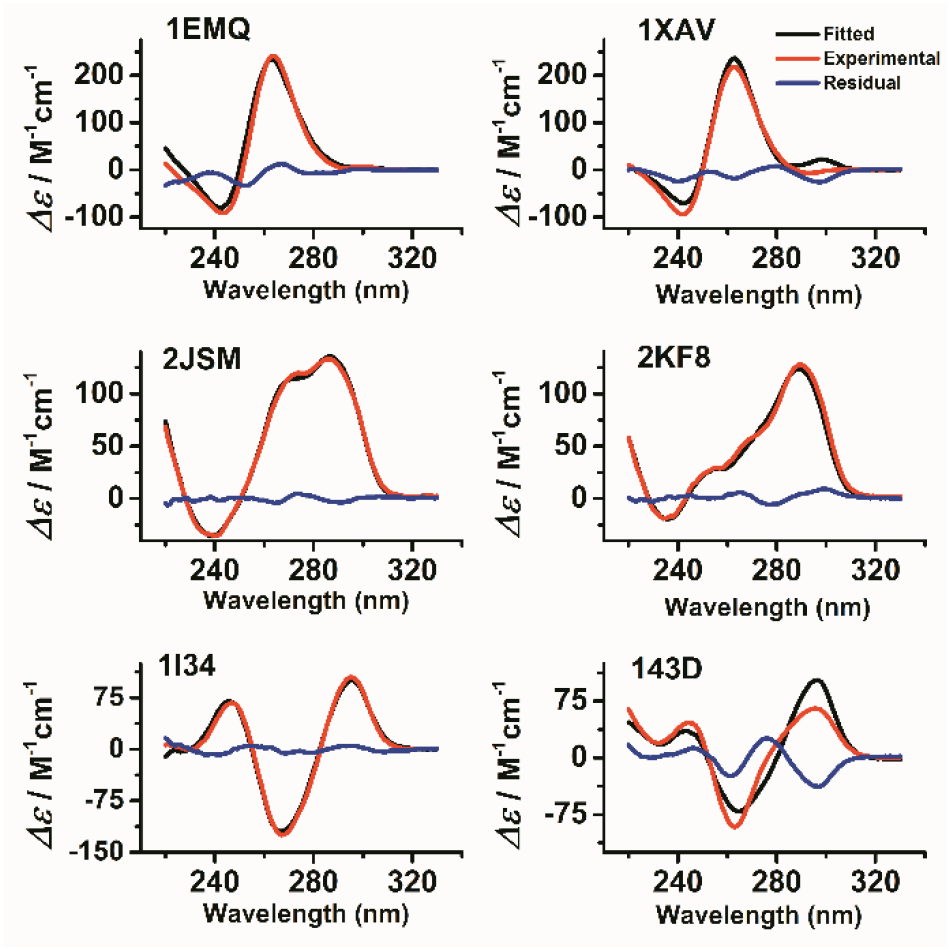
Experimental (red), fitted (black) and residual (blue) CD spectra for selected G-quadruplexes obtained by nonlinear least-squares fitting to eq. 1. The PDB identifier for each structure is shown in each panel.

To evaluate the accuracy of the prediction for each G-quadruplex CD spectrum, a leave-one-out cross-validation using different random initial guesses of secondary structural fractions was performed with a custom program (see supporting information). The estimated secondary structural fractions obtained by fitting (Table S5) are plotted against the known values (Table S4) determined from the deposited PDB structures in Figure 6. The correlations are excellent, indicating that our derived basis spectra can reliably be used to extract secondary structural content from a G4 CD spectrum.

**Figure 6.**
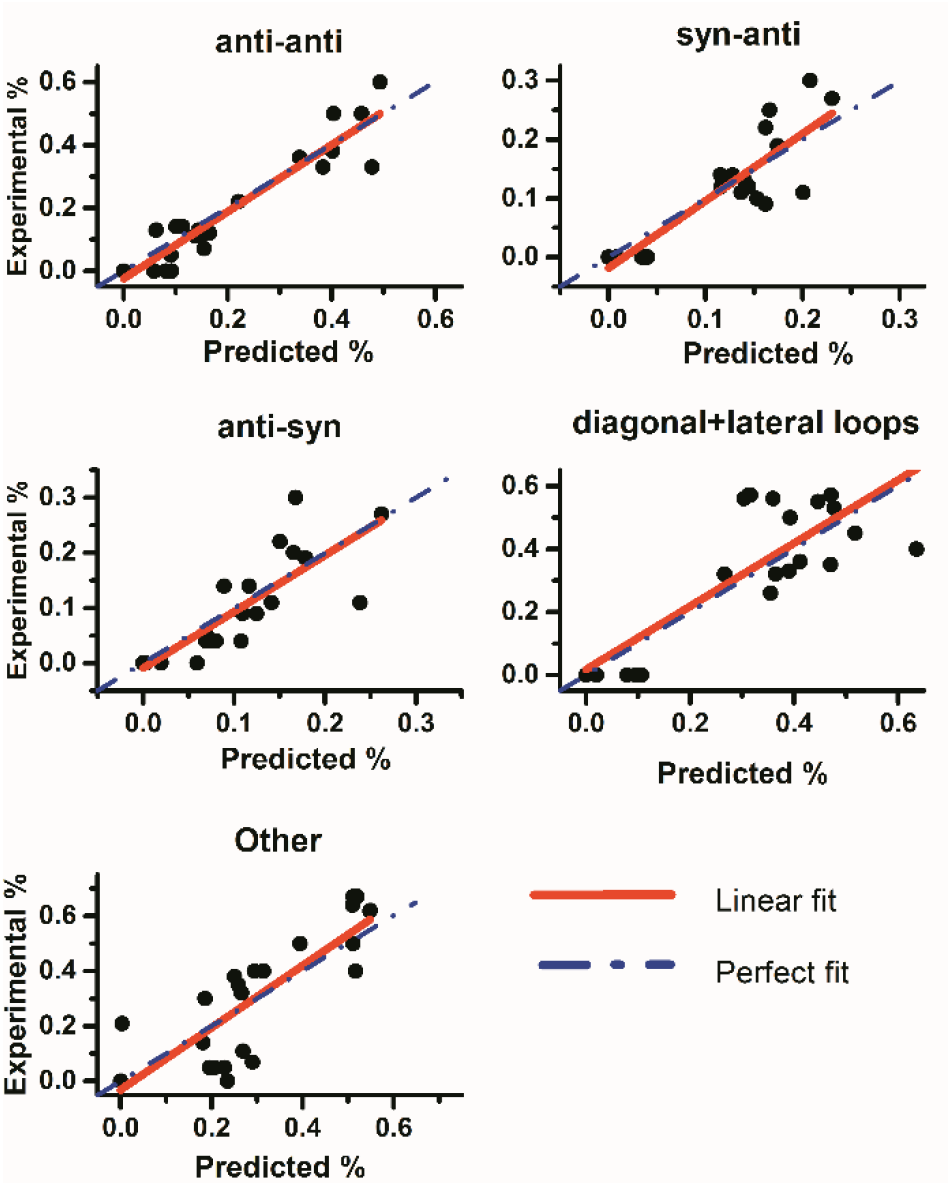
Scatter plots of the fractions of secondary structural elements determined from the known structures (“Experimental %”) versus the fractions obtained by fits to experimental CD spectra (“Predicted %). These data were obtained using the reference G-quadruplex CD spectra library and a leave-one-out cross-validation constrained least-squares fitting strategy. The red solid line is the least-squares linear fit to the data points, while the dashed blue line represents the line (slope =1) expected for a perfect correlation between the actual and estimated fractions.

The statistical analysis of the of the results for these structural parameters (Table S6) showed Pearson correlation coefficients and slope values closer to one, low structural RMSD, and values for the ξ parameter ^[13b]^higher than 1.6. These results demonstrate the accuracy of our method to predict the secondary structural parameters (*‘anti-anti’*, *‘syn-anti’*, *‘anti-syn’*, ‘diagonal and lateral loops’ and ‘other’) of a G-quadruplex by CD spectroscopy. In particular, the *‘anti-anti’* fraction was the secondary element that was most accurately predicted (Table S6).

Although CD spectroscopy cannot provide atomic level details of a G-quadruplex, the methodology described here represents a rapid and powerful tool to obtain quantitative secondary structural and topological information for a G-quadruplex in solution from its CD spectrum. This method can determine the secondary base step composition (*anti-anti, syn-anti, anti-syn, loops*) and the topologies (parallel, hybrid and anti-parallel) of G-quadruplex with accuracy. As far as we know, this is the first study showing that such detailed quantitative structural information of G-quadruplexes can be obtained by CD spectroscopy. This approach represents a significant advance for the characterization of G4 structures that complements higher-resolution NMR and crystallographic methods. Secondary structural information obtained by CD can be used to guide construction of molecular models for the structures of quadruplex-forming sequences in the absence of higher-resolution information.

The use of CD to determine protein secondary structure content evolved over several decades. Our study represents a first step in the development of an analogous CD tool for nucleic acids. As was the case for proteins, improvement and refinement of the CD method will be needed. For nucleic acids, this will require expansion of the reference library of known structures, construction of reference libraries with expanded wavelength spans that capture spectral features in the far UV, and algorithmic development to improve fitting procedures. We hope our study stimulates such efforts.

The fitting of G-quadruplex CD spectra described here was implemented in the open source R software environment (https://www.r-project.org/). Our script is available to interested users upon request (J.O.T.).

## Acknowledgements

Supported by grants CA35635 and GM077422.

